# Variability in cellular gene expression profiles and homeostatic regulation

**DOI:** 10.1101/021048

**Authors:** Yuriy Mishchenko

## Abstract

One of surprising recent discoveries in biology is that the gene and protein expression profiles in cells with identical genetic and environmental makeup can exhibit large variability. The nature and the significance of this variability had been posed as one of the current fundamental questions of biology. In this letter, we argue that the observed variability in cellular gene and protein expression can be understood as an outcome of homeostatic regulation mechanisms controlling the gene and protein expression profiles.

## INTRODUCTION

One of recent findings in biology is that the gene and protein expression profiles in cellular systems with identical genetic and environmental makeup can often exhibit large stochastic variability (*1–16*). The nature and the significance of this variability had been stated as one of the fundamental problems of biology. In this letter, we argue that the observed gene and protein expression variability can emerge naturally as the outcome of the regulation of gene and protein expression in biological cells by homeostatic regulation mechanisms.

One of the main challenges faced by biological systems is maintaining a stable physiological state in the face of the fluctuations in the parameters of internal and external environments. One of the best mechanisms available to biological systems for countering such disruptive influences is homeostatic regulation. Homeostasis is the property of biological systems to maintain a stable physiological state by means of various feedback mechanisms sensitive to the changes in that state (*17*). Homeostatic regulation responds directly to the changes of the controlled physiological parameters and, by doing so, can be very effective in maintaining the necessary values of those parameters. In this letter, we show that the property of homeostatic regulation to be sensitive directly to systems’ physiological states also leads with necessity to large variability observed in the internal configurations of such systems. This conclusion is general and is based broadly on two properties of biological systems – the possibility of implementing the same physiological state via different internal configurations and the reliance of homeostatic regulation on that physiological state for feedback. We present a general argument towards this point and demonstrate the emergence of this phenomenon in a simulated model.

## MATERIALS AND METHODS

We consider a simple model here whose goal is to inspect the impact of homeostatic regulation on the evolution of internal configuration states of a population of simple biological cells. Specifically, we consider a model of the control of a single protein expression in a cell, whereas (importantly) the protein concentration is regulated via several concurrent protein production pathways. The time-evolution of the protein concentration in such a cell is described by the following relationship, 

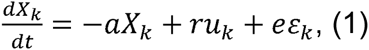
where *X_k_* is the concentration of the protein in the cell due to the regulation pathway *k*, *aX_k_* is the rate of the natural degradation of the protein, *ru_k_* is the rate of the protein production in pathway *k*, and *ε_k_* is the standard normal noise variable with zero mean and unit variance. *a,r,e* are constants and *k* is the index talking on the values from 1 to *N,* enumerating different protein production pathways, and *N* is the total number of such pathways. *u_k_* is understood as an internal control variable used by the cell to adjust the protein production rate in the specific pathway *k*. The key value of interest here is the total protein concentration in the cell given by 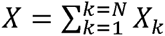

The model (1) is incomplete in an important way, and to complete this model it is necessary to determine how the control variables *u_k_* will depend on the cell’s internal state {*X_k_,k* = 1 *...N*}. We inspect three possibilities here: (i) feed-forward regulation, (ii) individual regulation, and (iii) homeostatic regulation. In feed-forward regulation, the values of *u_k_*are fixed at a constant value 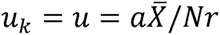, implying that the rate of protein production in each pathway is fixed in feed-forward manner, whereas 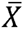 is the desired final protein concentration. The equilibrium is then achieved when *aX_k_* = *ru_k_* or 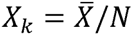 and, thus, 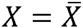. In the individual regulation, the pathways are regulated by the cell using individual feedbacks modeled by 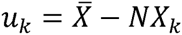. In this case, each pathway is driven towards a fixed point 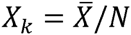 resulting in the same final protein concentration 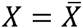. Finally, in the homeostatic regulation, the pathways are regulated via a common homeostatic feedback defined by 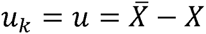 and controlled by the actual final concentration of the protein *X.* In this model, the regulating pathways are driven to a suitable configuration point by relying on the final protein concentrations in the cell.

## RESULTS

We argue that the large variability in the gene and protein expression levels observed in otherwise identical biological cells can be a direct consequence of homeostatic regulation mechanisms affecting the gene and protein networks of such cells. One of the main challenges faced by biological systems is the maintaining of their physiological state in the face of the fluctuations in the parameters of such systems’ internal and external environments. One of the best mechanisms available to biological systems for countering such disruptive influences is homeostatic regulation.

Homeostasis is the property of biological systems to maintain stable internal physiological state by means of feedbacks sensitive to the changes directly in such systems’ physiological parameters (*17*). By responding directly to the changes in physiological state, homeostatic mechanisms can be highly effective in maintaining the physiological conditions necessary for the normal functioning of biological organisms. When compared to other regulation strategies, homeostatic regulation can both achieve high specificity of physiological state’s regulation under heavy perturbations and noise as well as allow biological systems to recover from otherwise fatal failures, Figure 1.

**Figure 1:**
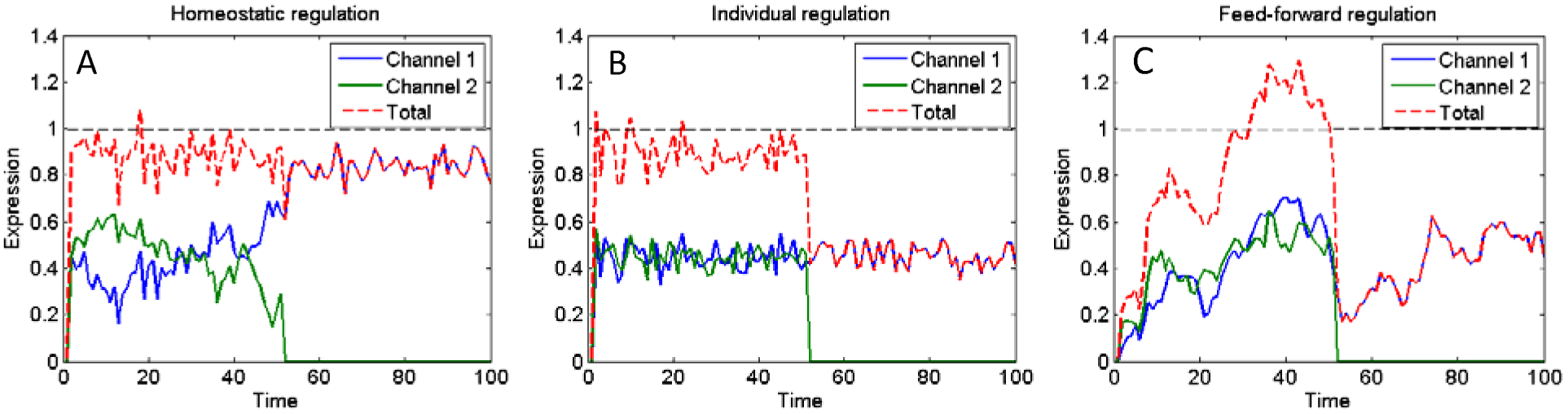
Time evolution of the total protein concentration in model of protein regulation in a cell, controlled by two complementary regulation pathways as discussed in the paper. A) Homeostatic regulation offers a high level of noise suppression in the final protein concentrations as well as allows recovering from failures in one of the pathways (such as pathway 2 blockage introduced here at time 50 units). B) Individual control of the protein production pathways, driving pathways towards a fixed operating point individually, also offers a high level of resistance to noise but fails to recover from the failure of one of the pathways. C) Simple feed-forward regulation both is highly vulnerable to noise and fails to recover from the failure in one of the pathways. Simulation parameters: *e* = 0.05, *a* = 0.1, *r* = 0.5. The target concentration 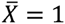 as indicated by the dashed line.

The capacity of homeostatic regulation to respond directly to the changes of controlled physiological parameters – the key element of homeostatic regulation responsible for the above important qualities – also makes homeostatic regulation insensitive to the perturbations that do not produce a significant change in a system’s physiological state. For example, these can be random up-regulations of one and simultaneous down-regulations of other pathways regulating the same physiological parameter. Whenever such a perturbation occurs, no net change may result in the physiological state causing no corrective actions from the homeostatic feedbacks.

To investigate this peculiar point more thoroughly, we turn our attention to a simple model of cellular regulation of a single protein’s concentration, *X,* controlled via two complementary production pathways, *X_1_* and *X_2_,* such that *X = X*_1_*+X*_2_. If the concentration of the protein in such cells’ cytoplasm is subject to a significant amount of noise, as can be expected in real biological cells due to the effects of spontaneous degradation of the protein as well as thermodynamic noise affecting the protein’s production, there can be multiple approaches for the cell to ensure that the concentration of the protein in the cytoplasm remains at the levels required for its normal functioning. One of the most basic such approaches is to maintain the rate of the protein production in each regulation pathway at a fixed level, whereas a given final concentration 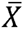 would be then achieved in the equilibrium depending on the protein’s overall production and degradation rates. Of course, such a simple “open-loop” regulation approach could be heavily affected by noise. A better approach may be to use a “closed-loop” feedback-regulation within each regulation pathway that ensures that the protein production in each pathway remains at specified levels, such as 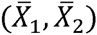 that result in required total protein concentration 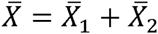. Yet another possibility involves regulating the protein production in all pathways using feedbacks relying directly on the protein’s total concentration in the cytoplasm, 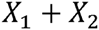, and actively enforcing 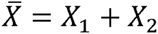 The latter, of course, is the homeostatic regulation. Of the three approaches, homeostatic regulation is the one that offers the highest degree of robustness to biological cells.

It is clear that, should we choose homeostatic regulation to control the protein concentration in our model, such regulation will be insensitive to the changes in the expression profiles of pathways *X*_1_ and *X*_2_ such that do not affect the protein’s total concentration *X* = *X*_1_+*X*_2_ This corresponds to up-regulating the protein production in one of the pathways while simultaneously down-regulating the other, so that the sum *X*_1_ + *X*_2_ remains intact. Obviously, it is always possible to produce such a perturbation and it is equally obvious that the latter homeostatic regulation would produce no action in the event of such a perturbation since the value of the total protein concentration *X* does not change. This insensitivity, however, can have dramatic consequences for the internal configuration states of such cells, (*X*_1_, *X*_2_) In Figure 2, we show the distribution of the internal states (*X*_1_, *X*_2_) of one such cell realized over different points of time. One can clearly see the spread of those states along the direction *X*_1_*+X*_2_ = *const,* where the homeostatic regulation provides no corrective feedback. This spread is caused by the accumulation of the above “neutral” perturbations of the cell’s state, that is, such perturbations that produce no net change in the total protein concentration *X* and, thus, cause no corrective actions from the homeostatic feedback. The accumulation of such “neutral” perturbations leads to a large variability of our model cells’ internal configuration states even when such cells otherwise share identical initial conditions and external environments.

**Figure 2:**
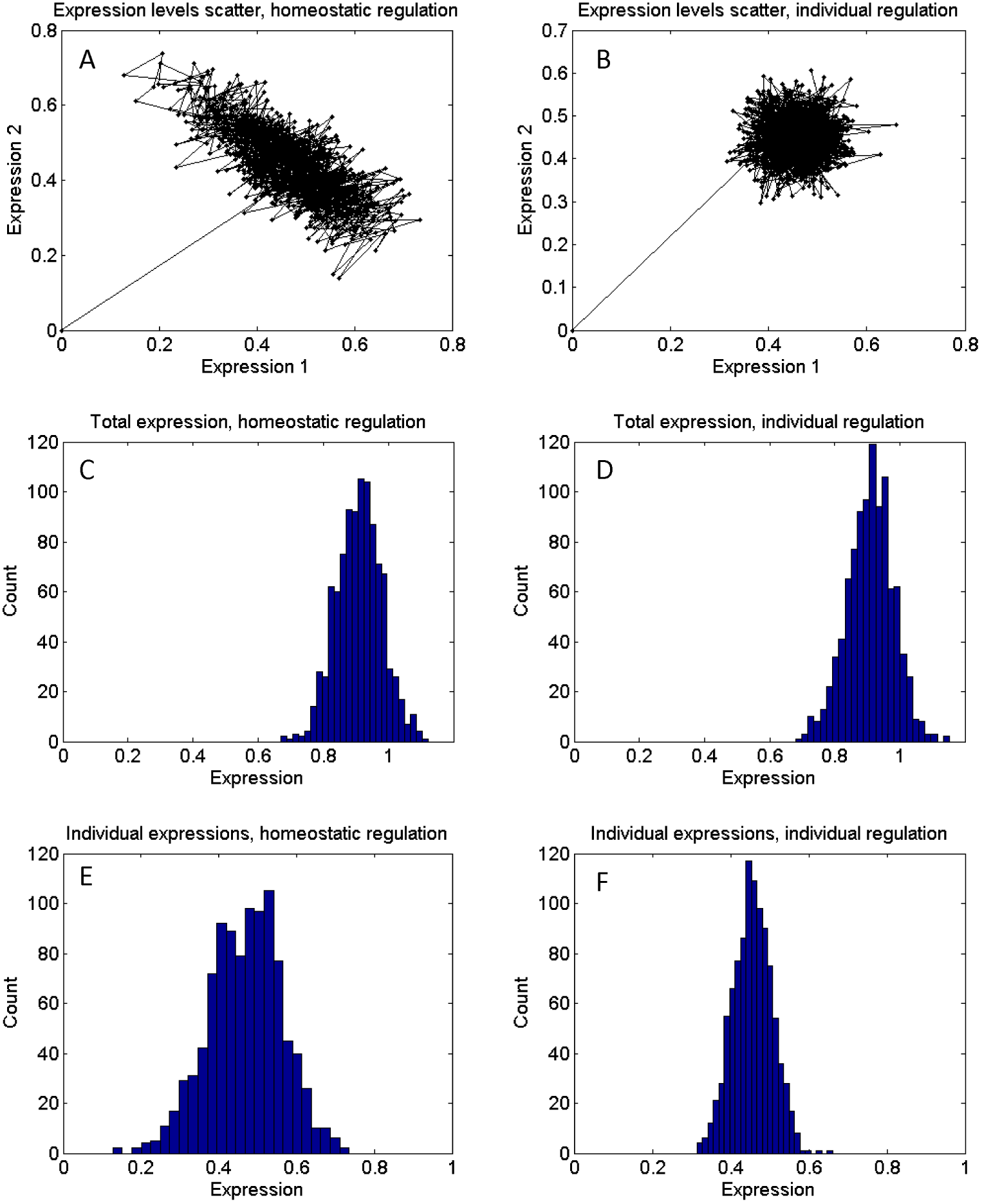
The insensitivity of homeostatic regulation to the perturbations of a biological system leaving the final physiological state unaffected results in a large spread of such systems’ internal states along the dimension of “insensitivity” of homeostatic regulation. A) Realized internal configuration states (*X*_1_,*X*_2_) in the model (1) under homeostatic regulation; the spread of the internal states along the direction of *X_1_+X_2_= X=const* is clearly visible. B) Realized internal configuration states for the model (1) with individually controlled pathways X_1_ and X_2_. C-D) In both examples, the degree of variation in the final protein concentration in either homeostatic regulation (C) or individual regulation (D) is the same. E-F) Under homeostatic regulation the spread in the expression levels of the individual pathways X_1_ and X_2_ (left, E) is at least two times larger than that under individual regulation (right, F).

The effect described above can become rather dramatic when the number of regulating pathways is large. In Figure 3, we show the example of the same model discussed above but now with *N* = 10 concurrent production pathways. In this case, the expression levels of individual protein production pathways *X_k_* vary widely, filling entire regions of configuration space and retaining no resemblance at all to the actually single physiological state that is realized in such model biological cell. The variation in the expression levels of individual protein production pathways here is dramatic and is up to 10 times greater than what would be expected from simple feed-forward or non-homeostatic feedback regulation, Figure 3E and 3F.

**Figure 3:**
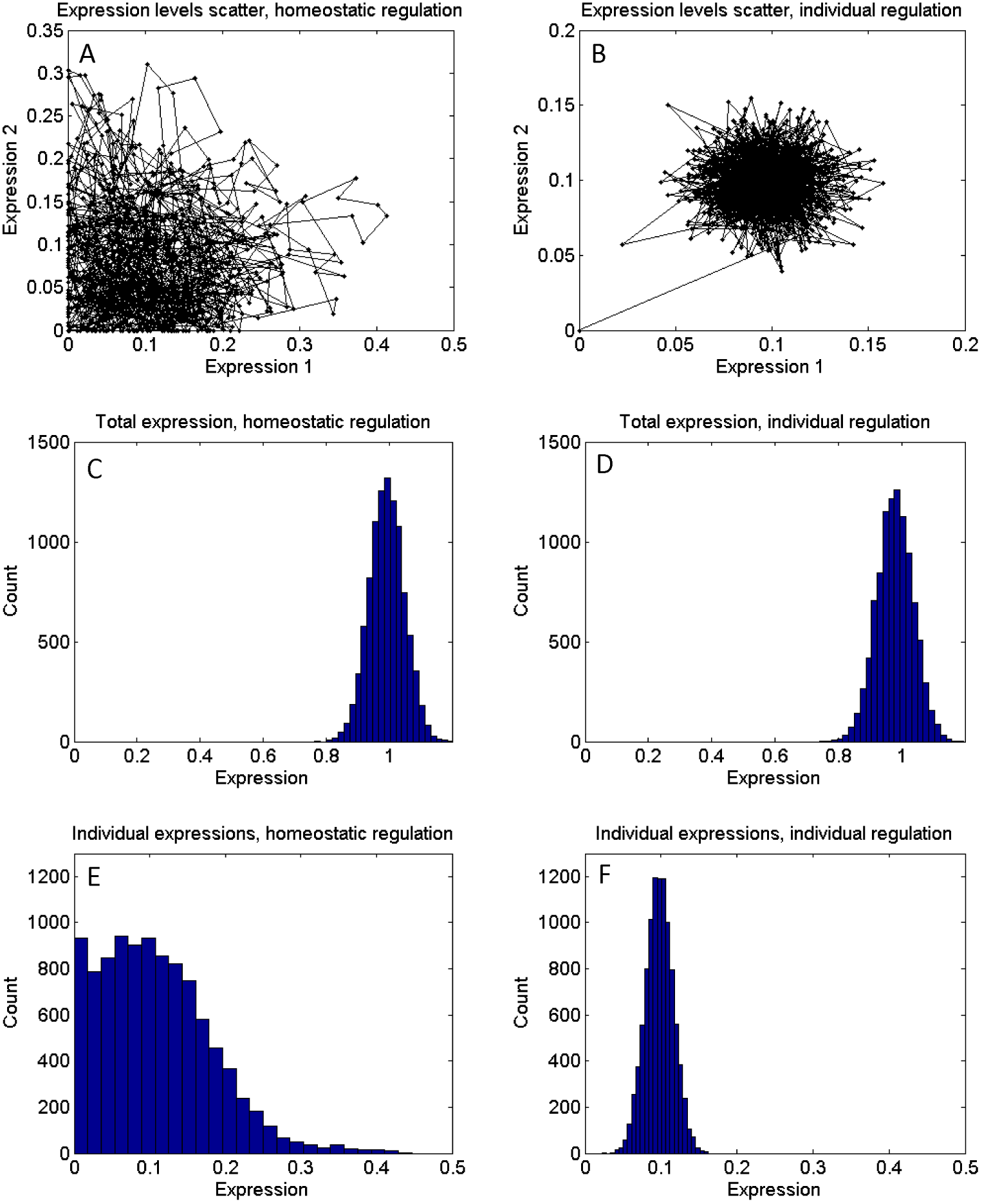
The stochastic variation in biological systems’ internal configurations may reach dramatic levels when the number of internal degrees of freedom, that is, the pathways regulating the same physiological parameter, is large. A) Realized internal states (*X*_1_,*X*_2_) of the model system (1) under homeostatic regulation with *N*=10 regulation pathways. B) Realized internal states for the model system (1) with *N*=10 under individually controlled pathways. C-D) The variation of the final total protein concentration is the same under either homeostatic (C) or individual regulation (D). E-F) The spread in the expression levels of the individual pathways *X_k_* under homeostatic regulation (E) is dramatically larger than that under individual regulation (F).

In conclusion, we find that the reliance of homeostatic regulation on biological systems’ actual physiological state can make such regulation insensitive to perturbations in such systems’ internal parameters that do not effect a net change in such systems’ physiological state. This makes it possible for such “neutral” perturbations to accumulate over time, resulting in large spread of such systems’ internal configuration states without any associated difference in such systems’ phenotype, genotype, or external environment.

## DISCUSSION AND CONCLUSIONS

We argue that the puzzling stochastic variability observed in the gene and protein expression of otherwise identical biological cells can be understood as a result of the homeostatic feedbacks in such cells’ internal homeostatic regulation mechanisms. Homeostatic regulation is ubiquitous in biology. Compared to other regulation mechanisms, homeostatic regulation offers superior capacity towards reducing stochastic fluctuations in physiological state as well as the ability to recover from otherwise critical failures. The key property of homeostatic regulation lies in its responding to the changes of the actual physiological state of a system. This gives homeostatic regulation superior ability for countering disruptions of physiological state but also makes it insensitive to biological systems’ perturbations that do not produce a net change in the values of controlled physiological parameters. Accumulation of such “neutral” perturbation can lead to large variations in the internal configurations of biological systems that are otherwise identical.

The wide spread of homeostatic regulation in biology implies that this phenomenon is likely to emerge not only in the gene and protein expression networks in biological cells, but also much more widely as a generic property of biological systems. Indeed, we note other possible examples of the same phenomenon that had been reported in the literature with respect to the axonal and dendritic ion-channel composition in neurons and the structure of central pattern generator neural circuits in lobster somatogastric ganglion (*17–22*). One can expect more similar examples to emerge in the future.

If homeostatic regulation is the primary cause of the stochastic variability observed in cellular gene and protein expression in biological cells, then the discussion above allows us to make certain experimentally verifiable predictions. For instance, we can expect that higher amounts of such variability will be reported in the systems with a larger number of internal degrees of freedom, that is, where more complementary pathways can contribute to the regulation of the same physiological parameter. Furthermore, such stochastic variability should be completely decoupled from the variability of the phenotype, while the variations of individual gene and protein expression levels should be, on the contrary, quite strongly correlated. More interesting is the prediction that the gene and protein expression profiles within individual cells should change over time. That is, the said stochastic variability can be observed not only over a population of similar biological cells, but also in a single cell where the gene and protein expression profiles are measured at different points of time. Finally, we expect similar examples of stochastic variability to be found in many other biological systems ranging from single cells and multicellular organisms to complex biological systems such as neuronal circuits.

## Acknowledgement

This work had been supported by the Bilim Akademesi-The Science Academy under the BAGEP program (Turkey), Toros University BAP grant number TUBAP135001 (Turkey), and TUBITAK ARDEB 1001 grant number 113E611 (Turkey).

